# Hyperaccumulation of Vanadium in animals: two sponges compete with urochordates

**DOI:** 10.1101/2023.06.02.543482

**Authors:** Kassandra de Pao Mendonca, Perrine Chaurand, Andrea Campos, Bernard Angeletti, Mauro Rovezzi, Ludovic Delage, Carole Borchiellini, André Le Bivic, Julien Issartel, Emmanuelle Renard, Clément Levard

## Abstract

Vanadium (V) is the second most abundant transition metal in the oceans. Nevertheless, its concentration in organisms is generally low despite its involvement in various physiological or enzymatic functions. Because exposure to high concentrations was shown to be harmful for various organisms including animals, the unexpected very high concentrations found in urochordates and annelids yielded additional studies to decipher their biological meaning.

Here we report cases of V hyperaccumulators in a distinct animal phylum (Porifera): the two homoscleromorph sponge species *Oscarella lobularis* and *Oscarella tuberculata* (up to 30 mg/g dw). These high concentrations overpass those reported previously in urochordates and annelids and are not found in the 5 other sponge species studied here. In both *Oscarella* species, vanadium is mainly accumulated in the 100 μm surface tissues, and in particular in mesohylar cells, as vanadyl (+4) before being partly reduced to V (+3) in the deeper tissues. Genomic surveys failed to find any previously described gene implicated in vanadium metabolism, suggesting that V hyperaccumulation emerged convergently in this lineage. This feature may be of interest for developing bioremediation strategies in marine ecosystems or bioinspired processes to recycle this critical metal.

**Synopsis:** This study reports extremely high concentrations of vanadium in two sponges. This unusual bioaccumulation of reduced V (+3) in mesohylar cells relies on unknown molecular mechanisms.

## Introduction

Vanadium (V atomic number 23) is one of the most abundant elements (21^st^) on Earth and represents the second most abundant transition metal in oceans (Awan et al., 2021; Jeandel et al., 1987; Morford & Emerson, 2018; Schlesinger et al., 2017). It is naturally present in the terrestrial crust, river waters, and atmosphere (Huang et al., 2015; Schlesinger et al., 2017) and can show local enrichments, as for example in volcanic areas (Rehder, 2015). In nature, this redox sensitive element exists in three oxidation state forms: V (+3), V (+4) (i.e., vanadyl), and V (+5) (i.e., vanadate), and in a variety of coordination environments (from octahedral (Oh) to a five-coordinate (Py) or a tetrahedral geometry (Td)), depending on redox conditions and the pH of the surrounding environment (Wehrli & Stumm, 1989). In sea waters, the average V concentration is between 1 and 2 µg/L and Vanadium is often found as the vanadate form H_2_V^5+^O_4_^−^ (Nielsen, 2020).

Despite this abundance, Vanadium is considered as one of the 30 critical metals and materials listed by the European Union in 2020 (EU, 2020). V is used in a large variety of applications such as the manufacture of chemical catalysts, high-strength low-alloy steels for the aerospace and aviation industries as well as in nuclear reactors (Yan et al., 2021). It is also involved in the production of pesticides, dyes employed in textiles and ceramics and in next-generation flow batteries (Moskalyk & Alfantazi, 2003; Treviño et al., 2019). During the last decade, V has also been extensively studied for medical applications, especially in the formulation of treatments against diabetes and cancer (Treviño et al., 2019). The increasing use of V and of its subsequent release in the environment represent a threat for ecosystems (Moskalyk & Alfantazi, 2003; Schlesinger et al., 2017; Watt et al., 2018). For example, high quantities of V are released into the atmosphere *via* the combustion of fossil fuels (Schlesinger et al., 2017; Zoller et al., 1973) or from V-derived products stored in landfills (Larsson et al., 2015; Power et al., 2011). Finally, anthropic V is transported into the oceans *via* atmospheric fallout, rivers, and effluents; this means that all continental and marine ecosystems are concerned (Tulcan et al., 2021).

In contrast with other transition elements (Mo, W, Mn, Fe, Co, Ni, Cu and Zn), V does not seem to be an essential bioelement for most organisms (Gustafsson, 2019). Indeed, V-dependent biological functions or enzymes (vanadium nitrogenases and vanadate-dependent haloperoxidases) have so far been reported only in a limited number of organisms (Rehder, 2015). This metal is thus often present in low amounts in most organisms with concentrations generally in the ng/g dry weight (dw) range, rarely up to µg/g dw (Fukushima et al., 2009). Exposure to relatively high concentrations of V (in the mg/L range) was shown to be toxic in most organisms, such as algae, land plants, and freshwater planktonic species (Wang & Liu, 1999, Schiffer & Liber, 2017, Gustafsson, 2019). This toxicity is mainly explained by the high similarity between vanadate and phosphate. Vanadate can thereby substitute for phosphate in enzymes such as phosphatases, kinases, phosphomutases,diesterases, ATPases and ribonucleases, thereby resulting in the inhibition of these enzymes (Huyer et al., 1997; Rehder, 2015; K. H. Thompson et al., 2009).

Considering this toxicity, the accumulation of very high amounts of V in a few species seems paradoxical. Among Metazoa (animals), the most well-known hyperaccumulators of V are urochordates Ascidiacea (sea squirts) and annelids of the Sabellidae family (fan worms) (E. D. Thompson et al., 2018). In fan worms, V concentrations range from 90 to 10,500 μg/g dw depending on the considered species (Fattorini et al., 2010; Fattorini & Regoli, 2012; Ishii et al., 1994; Popham & D’Auria, 1982). In ascidians, where V body levels vary among taxa, Phlebobranchia suborder shows higher levels of V than that of the Stolidobranchia suborder. For example, in *Ciona edwardsi, Ascidia ahodori, Phallusia mammillata* and *Ascidia gemmata*, V concentrations in “blood” cells (named signet ring cells or vanadocytes) range from 1,500 to 21,000 µg/g dw which represents an enrichment factor from seawater of 10^4^ (Michibata et al., 1987, 1991). The proteins involved in the transport, storage, and metabolism of V in this ascidian lineage was thoroughly studied (Kawakami et al., 2009; Ueki & Michibata, 2011) but its biological role is still a matter of debate (Fattorini & Regoli, 2012; Ueki & Michibata, 2011).

The present study reports for the first time hyperaccumulation of V in two sponge species (Porifera) from the Mediterranean Sea. V distribution at the tissue and cell scales was determined and the oxidation state of V was investigated. Finally, we performed a gene survey to decipher whether some molecular mechanisms are shared with urochordates and annelids.

## Materials and methods

### Sampling

Seven sponge species (Porifera) from 3 different classes were sampled (Fig.1A) in the Mediterranean Sea by divers: one Calcarea (*Clathrina clathrus* (Schmidt, 1864)), three Demospongiae (*Agelas oroides* (Schmidt, 1864), *Aplysina cavernicola* (Vacelet, 1959) and *Axinella damicornis* (Esper, 1794)) and three Homoscleromorpha (*Corticium candelabrum* (Schmidt, 1862), *Oscarella tuberculata* (Schmidt, 1868) and *Oscarella lobularis* (Schmidt, 1862)). All species (at least 3 individuals/species) as well as one sample of *Ciona edwardsi* (Urochordata, for comparison) were collected at the same site, Maïre island (43°12.760’N, 5°19.800’E) in the bay of Marseille (South littoral zone of France) where they can be found in sympatry (in contrast, Hexactinellida are absent at this sampling site). Concerning *O. lobularis* and *O. tuberculata*, additional samples were collected at other sites, depending on the distribution of the species: 6 other sites in the bay of Marseille and one site in the bay of Toulon. The Toulon sampling site was selected from a strategic point of view since Toulon is subjected to many industrial, military and port activities (Tessier et al., 2011). In addition, in order to evaluate environmental concentrations, surface sediments were manually collected at each of the eight sites (Supporting Information 1).

**Figure 1:**
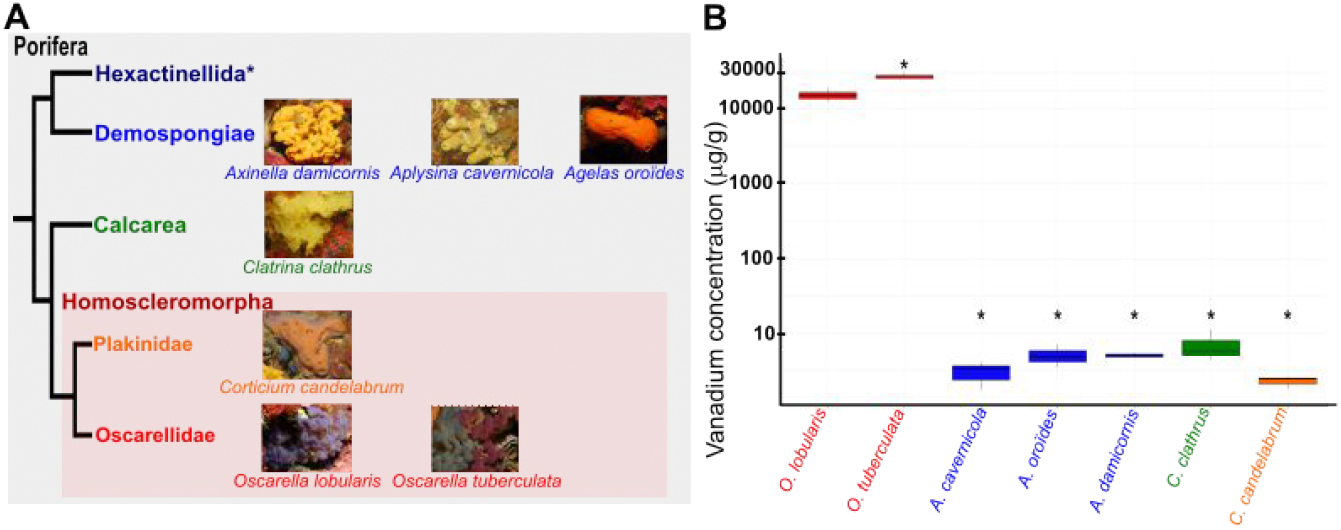
(A) Phylogenetic relationships between the four classes of Porifera and phylogenetic position of the sampled species. Hexactinellida (*) are found only in deep seas or marine caves and therefore cannot be found at the Maïre site. (B) Boxplot of V concentrations in sponge species sampled at the Maïre site (µg/g dw). The color code used for sponge classes is the same as in A. The stars show a significant difference between the considered sponge and O. lobularis (Mann–Whitney U test with p <0.05).

### Vanadium concentrations in marine sponges and sediments

Samples were transported, cleaned and frozen exactly as described in detail in de Pao Mendonca et al. (2023). Then, 300 mg dw of each sponge or 50 mg dw of sediments were digested by acid treatment and mineralized with a microwave (see de Pao Mendonca et al., 2023, for a detailed protocol). After mineralization, the samples were diluted by a factor of 5-100 before analysis by Inductively Coupled Plasma Mass Spectrometry (ICP-MS Perkin Elmer, NexION® 300X). For each analysis series, technical replicates, procedural blank, and a standard sample (corn powder V463, INRAE) were analysed (Fehlauer et al., 2022) (Supporting Information 1). Results are expressed as µg/g of V dw and represent mean values obtained for each site.

### Cellular localization of vanadium in Oscarellidae species

An adult sponge of each of the two sister species *O. lobularis* and *O. tuberculata* was collected at the Maïre site and then transported and cleaned as previously described. They were fixed using glutaraldehyde 2.5 % in a mixture of 0.4 M cacodylate buffer and seawater as described in Ereskovsky *et al*., 2013 but without osmium (Ereskovsky et al., 2013). A cryofracture with liquid nitrogen was performed to prevent damaging internal structures. Finally, the samples were observed by Field Emission Gun Scanning Electron Microscopy (SEM, Zeiss GeminiSEM 500) coupled with an Energy Dispersive x-ray (EDAX SDD) detector, at 20 kV. 30 spots per cell type were analysed on the first 200 micrometres of the sponge tissue in order to determine the concentration of V for each cell type. Simulations performed with “Monte Carlo Simulation of electron trajectory in solids” (CASINO) V2.4 software, reveal that during EDX analysis of the marine sponges, the incident electron beam penetrates the area of ∼ 1.5 µm depth for V (Supporting Information 2). The size of the electron beam interaction is therefore compatible with the size of the cells targeted in the sample (smaller cells are about 7µm long).

### Oxidation state of vanadium measured in-situ in Oscarellidae species

V speciation in *O. lobularis* and *O. tuberculata* species sampled at the Maïre site was determined by X-ray absorption Spectroscopy (XAS) at the European Synchrotron Radiation Facility (ESRF, Grenoble, France). For comparison, the V speciation was also determined in the sample of *Ciona edwardsi* sampled at the same site. Prior to analysis, the organisms were transported and cleaned as previously described and stored at -20° C. For the analysis, samples were cut into 1 mm-thick cross sections in liquid nitrogen using a scalpel and then directly transferred to the XAS sample holder.

High Energy Resolution Fluorescence Detected X-ray Absorption Near Edge Structure (HERFD-XANES) spectra were collected at the V K-edge (5465 eV) on the FAME-UHD beamline (French CRG at the ESRF) (Levard et al., 2025). The incoming X-ray energy was monochromatized using a Si (220) cryogenically cooled and sagittally focusing double crystal monochromator. The monochromator was calibrated using a V metallic foil. The first maximum of the spectrum first derivative of the X-ray absorption spectrum was set at 5465 eV. The spectra were collected in step scanning mode with a beam size of approximatively 0.2 × 0.1 mm^2^. The sample temperature was kept at 10 K using a liquid Helium flow cryostat to prevent sample beam damage. The relatively small beam height (0.1 mm) is adapted to discriminate the average V speciation from surface tissue (100 µm thick) to internal tissue within the sponge cross-section. Samples were analysed in fluorescence mode using a crystal analyser spectrometer equipped with 10 Ge (331) spherically bent crystal analysers of 1 m radius of curvature aligned at the V Kα1 line (4953 eV, Bragg angle of 74.63°). The measured experimental broadening of the spectrometer was 1.6(1) eV. One spectrum represented an average of 4-6 scans for reference compounds, 2-7 scans for sponge tissues and 3 scans for ascidian.

A set of reference compounds in which V occurs in the three oxidations states (+3, +4 or +5) in coordination geometry from octahedral (Oh) to a five-coordinate (Py) or a tetrahedral geometry (Td) was prepared at 4000 ppm of V to avoid self-absorption issues. Reference compounds were either diluted in boron nitride (BN) and pressed into pellets for solids or dispersed in ultrapure water for certain salts. It includes commercial solids VCl_3_, V_2_O_4_, VOSO_4_, nH_2_O, vanadyl acetylacetonate C_10_H_14_O_5_V, V_2_O_5_, Na_3_VO_4_ purchased from Aldrich, a solution of Na_3_VO_4_ dispersed in ultrapure water and a natural mineral Pascoite Ca_3_(V_10_O_28_).17H_2_O. Oxidation state and symmetry of V in each of the reference compounds are detailed in Supporting Information 3 and 4. Finally, calibration, normalization, and data averaging were performed using Larch XAS Viewer 0.9.65 software.

Pre-edge analysis was carried out following a procedure inspired by Chaurand et al. (2007) and Levina et al. (2014). They demonstrated that the pre-edge features can be used for the determination of average speciation of V in complex biological or environmental matrices. They reported noticeable changes in both intensity and position of the pre-edge peak according to V oxidation state and coordination number, using a range of reference compounds. The pre-edge features were obtained by fitting a baseline-corrected pre-edge peak with a series of mixed Gaussian-Lorentzian curves (pseudo-Voigt functions) (Supporting Information 4). Care was taken in using the smallest possible number of components. The “pre-edge peak area, PEA” (sum of the integrated areas of each component normalized to V_2_O_5_ pre-edge area) and the “pre-edge peak centroid energy, PECE” (area-weighted average of the position in energy of each component) were calculated.

### Survey of genes involved in vanadium metabolism

Several genes involved in V transport, storage or metabolism in different metazoan species were searched for by blastP against the predicted proteome of *Oscarella lobularis* (genome draft (Belahbib et al., 2018)) using the local blast tool of Bioedit (Hall, 1999). The returned hit-blast with e-value < 10^−2^ were then checked by a reciprocal best-hit approach against the NR database (NCBI), PDB and Swiss-Prot for domain predictions. The list of query sequences and of best hits obtained are provided in Supporting Information 5. The same strategy was adopted using, as queries, bacterial genes involved in V metabolism on the predicted proteome of *Candidatus Rhodobacter lobularis*, the dominant microorganism associated with this sponge species (Jourda et al., 2015). The list of query sequences and of best hits obtained are provided in Supporting Information 6. In order to refine the assignment of candidate sequences, phylogenetic analyses were performed. Alignments (available upon request) were performed with MAFFT (Kuraku et al., 2013) followed by manual JALVIEW curation (Waterhouse et al., 2009). Phylogenetic analyses were performed using PHYML (WAG+G+I SH-LIKE model) (Guindon et al., 2010) and the trees were visualized with iTOL Newick tree (0.8-1 bootstrap cutoff) (Letunic & Bork, 2021).

### Statistical analysis

In order to evaluate the statistical significance in V sponge body contents, a non-parametric Kruskal-Wallis test was used (significant p-value < 0.05) for comparisons between sites, and a non-parametric Mann–Whitney U test was performed (p-value <0.05) to compare V concentrations between species. To analyze the significant differences in V quantities between cell types, a Mann–Whitney U test was used with a p-value <0.05. All statistical analyses and graphics were performed using R studio open-source edition package (https://www.rstudio.com/).

## Results and Discussion

### Vanadium hyperaccumulation in Porifera is a specificity of Oscarellidae

Seven sponge species, members of three of the four classes of the phylum Porifera were sampled at the same site (Maïre island) where they live in sympatry and were analyzed by ICP-MS (Fig 1A; Supporting Information 1). Five of them, the three Demospongiae (*A. cavernicola, A. oroides, A. damicornis)*, the Calcarea *C. clathrus* and the Homoscleromorpha Plakinidae *C. candelabrum* exhibit relatively low concentrations of V in their tissues ranging from 2.3 to 7.21µg/g dw (Fig.1B). Slight variations (not statistically significant) are observed between these five species: *C. candelabrum* and *A. cavernicola* harbor the lowest concentrations (2.30 ± 0.43 µg/g and 3.06 ± 1.21 µg/g dw respectively). In contrast, *C. clathrus* contains the highest vanadium concentrations (7.21 ± 3.65 µg/g dw). These values are lower than those found in the upper sediments collected at the same site (Maïre: 28.4µg/g dw, Supporting Information 1), suggesting that these species do not bioaccumulate V (*i*.*e*., exhibiting a bioconcentration factor (BCF)<1, Supporting Information 7). Unexpectedly, the two remaining species *Oscarella tuberculata* and *O. lobularis*, both members of Homoscleromorpha Oscarellidae, contain significantly higher concentrations of Vanadium - 2,000 to 15,000 times higher-than any other sponge species studied here (Fig.1B). These vanadium concentrations are also higher than the concentrations previously reported in 8 other demosponges ranging from 2 to 76.9µg/g dw in the Mediterranean Sea (Cebrian et al., 2003; Perez et al., 2005; Pérez et al., 2004). The highest V concentrations are found in *O. tuberculata* with values ranging from 25,198 to 30,274 µg/g dw. In the sister species, *O. lobularis* concentrations range from 12,411 to 18,425 µg/g dw. To the best of our knowledge, such concentrations of V have never been reported before for marine sponge species. The BCF of these two species is higher than 1 (Supporting Information 7). It is worth noting that the concentrations measured in *O. tuberculata* exceed those of C*iona edwardsi*, sampled at the same site (235 µg/g dw; Supporting Information 1), and those previously reported in urochordate species (Michibata et al., 1987, 1991). The hyperaccumulation of V by the two Oscarellidae species reported here is therefore of great interest.

### The hyperaccumulation of vanadium is not specific to the Maïre site populations

In order to test whether the hyperaccumulation of V was a specificity of Oscarellidae individuals growing at the Maïre site, we collected additional specimens of *O. tuberculata* and *O. lobularis* at 6 other sites in the bay of Marseille and 1 site near Toulon (Fig. 2A). Even if statistically significant variations between sites exist, high V concentrations are observed in Oscarellidae whatever the site. For the sites where they are found in sympatry, *O. tuberculata* always shows higher V concentrations than *O. lobularis*, with values ranging from 25,198 to 33,479 µg/g dw (Fig.2B). In addition, the adults collected at the previously mentioned Maïre site are, in fact, the specimens with the lowest concentrations in V with 15,204 ± 2,011 µg/g dw in *O. lobularis* (p value <0.05) and with 27,009 ± 2,833 µg/g dw in *O. tuberculata*. In contrast, the samples collected at the Fourmigues site, located near Toulon, show the highest concentrations in *O. lobularis* (28,212 ± 6,457 µg/g dw) followed by Riou site in the Marseille Bay (19,257 ± 9,508 µg/g dw).

**Figure 2:**
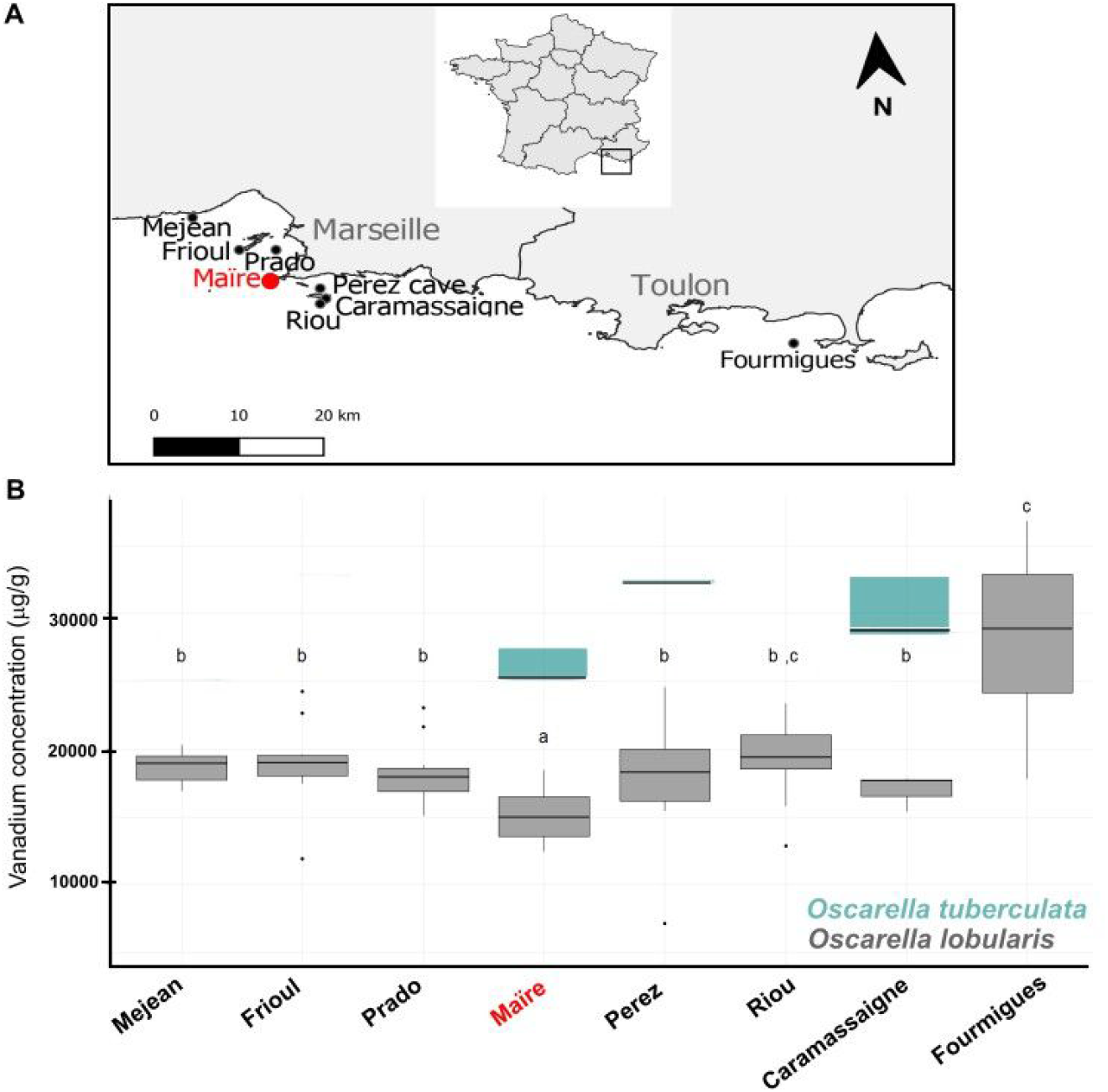
(A) Location of the sampling sites (Southern of France; North-Western Mediterranean Sea). According to the geographic distribution of Oscarella lobularis populations, 7 sites in the bay of Marseille (Mejean, Frioul, Prado, Maïre, Perez cave, Caramassaigne, Riou) and 1 site out of the bay (Fourmigues) were sampled. The red dot represents Maïre Island (Fig. 1), (B) Boxplot of V concentrations in O. lobularis and O. tuberculata according to the different sampling sites. In the boxplot, a different letter indicates a significative difference (p <0.05).

### Vanadium concentrations are higher in the mesohylar cells

A sponge’s body plan is very different from that of the other V-hyperaccumulative animals belonging to urochordates and annelids. Even though sponges are sessile filter feeders like urochordates, they are devoid of organs, vascular system and blood cells. In sponges, filtration is performed *via* a completely different system (unrelated to a digestive tube) named the aquiferous system (Borchiellini et al., 2021) (Fig.3A and Supporting Information 8). For this reason, it is necessary to determine in which part of the tissues this accumulation takes place and which cell types are involved.

**Figure 3:**
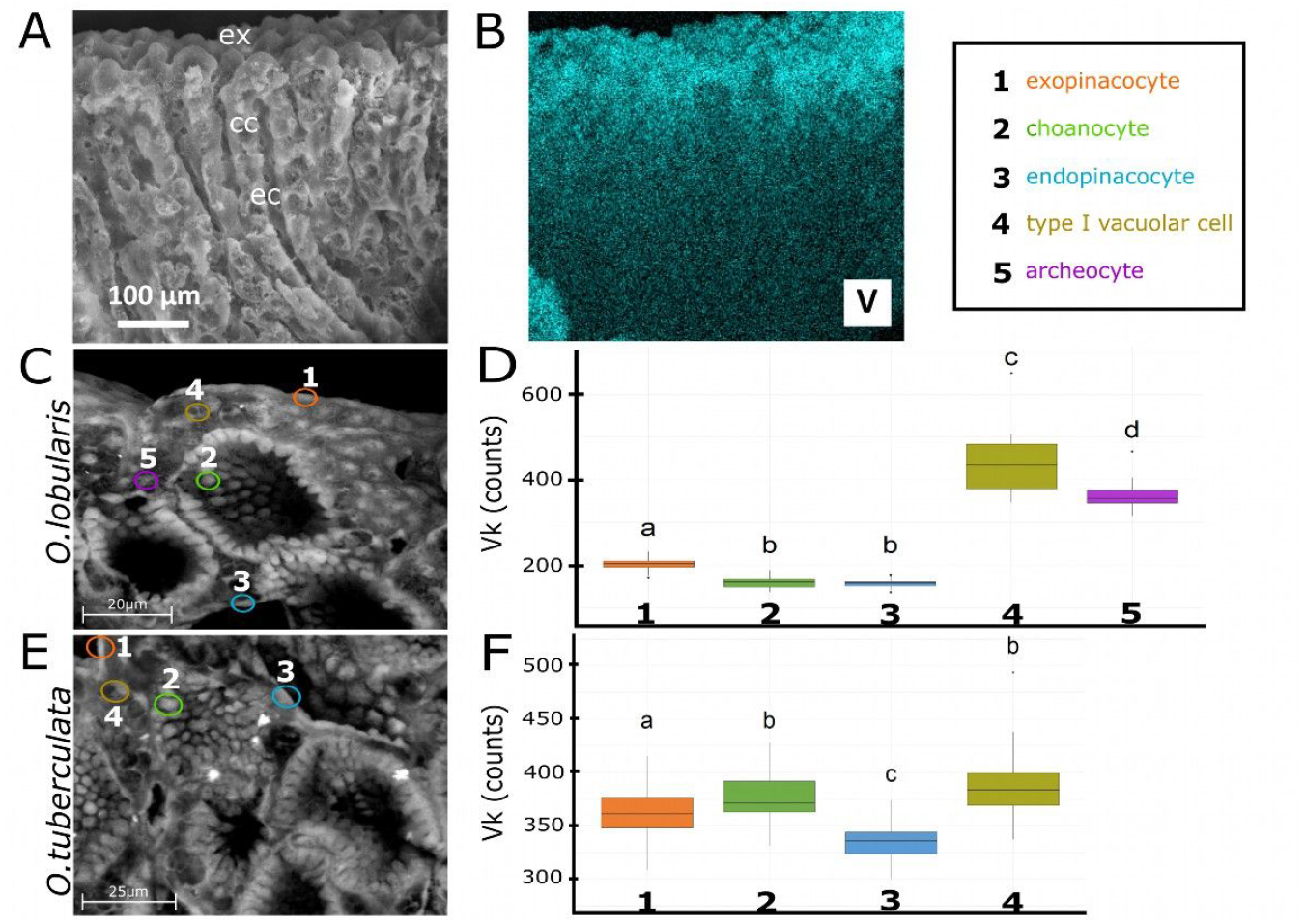
Localization of V in the tissues and cells of Oscarellidae species. (A) Backscattered SEM image of O. lobularis tissues (cryo section). ex: exopinacoderm, ec: excurrent canal (bordered by endopinacocytes), cc: choanocyte chambers. (B) V chemical distribution in O. lobularis from EDX analysis mapping (Kα emission line). (C, E) SEM images of the cellular organization in (C) O. lobularis and in (E) O. tuberculata. The only difference between O. lobularis and O. tuberculata is the presence of only one vacuolar cell type (type I vacuolar cell) in the mesohyl. (D, F) Vk quantities (counts) from microanalyses performed with electron beam pointed in the different cell types present in (D) O. lobularis tissues and in (F) O. tuberculata (N= 30 for each cell type). In the boxplot, a different letter indicates a significative difference (p <0.05).

In *O. lobularis*, the energy dispersive X-ray (EDX) mapping analysis shows a concentration gradient from the sponge surface down to the inner tissue (Fig.3B): indeed, V (and P) concentrations are higher in the 100μm surface tissues, in contrast to other elements such as C, S or K for which the distribution seems homogeneous (Supporting Information 9).

At higher magnification, the different cells can be clearly identified (Fig. 3C and 3E): the outer epithelium called the exopinacoderm (ex) is composed of exopinacocytes (1). The choanocyte chambers (cc) are composed of choanocytes (2). The canals of the filtration system (ec) are lined by endopinacocytes (3). Between these epithelium layers, the mesohyl is a layer composed of vacuolar cells and bacteria (Supporting Information 8). The separate analysis of chemical elements in the different cell types (4 for *O. tuberculata*, 5 for *O. lobularis*) (Fig.3C and 3E) shows that in both species the mesohylar cells (type I vacuolar cells and archeocytes (also named type II vacuolar cells)) have a significantly higher V concentration than exopinacocytes, endopinacocytes or choanocytes. Among mesohylar cells in *O. lobularis*, type I vacuolar cells display a higher V concentration than archeocytes with 444 ± 78 Vk counts. According to the statistical analysis, both choanocytes and endopinacocytes show significantly the lowest concentrations of vanadium (160 ± 13 Vk counts and 154 ± 16 Vk counts respectively) (Fig.3D). Regarding *O. tuberculata*, type I vacuolar cells and choanocytes present the highest concentration of vanadium (386 ± 30 Vk counts and 376 ± 23 Vk counts respectively); while endopinacocytes present a lower vanadium concentration with 334 ± 21 Vk counts (Fig.3F).

The fact that, in both species, the highest concentrations of vanadium are found in type I vacuolar cells, provides new insights into the largely unknown functions of this cell type, even though a storage/secretion property was already suggested, because of the presence of two to four large vacuoles (Gaino et al., 1986). Interestingly, in the V hyperaccumulator annelid *P. ocellata*, high levels of vanadium have been detected in the epidermal cells of the gill crowns within the apical vacuoles (Ishii et al., 1994). In ascidians, vanadium is also stored in the vacuoles of vanadocytes (Botte et al., 1979; Kalk, 1963).

### Vanadium is mainly reduced to vanadyl when absorbed by the Oscarellidae species

HERFD-XANES spectra obtained for sponges (Oscarellidae species) and ascidian (*Ciona edwardsi*), collected at the Maïre site are shown in Supporting Information 3. Pre-edge fitting is detailed in the Supporting Information 4 and 10. The plot of pre-edge features comparing model compounds and samples (PEA *vs*. PECE, Fig 4.B) reveals that V is incorporated by the Oscarellidae species mainly as vanadyl in both surface and internal tissues (V (+4)). Indeed, pre-edge features (PECE and PEA) are similar to those of reference compounds in which V occurs as V (+4) in octahedral geometry (Oh), especially VOSO_4_, nH_2_O (Supporting Information 10). Vanadyl is stable in moderately reducing environments, particularly when the organic matter concentration is high. It is highly reactive and forms a large number of strong complexes with organic ligands (Gustafsson, 2019).

**Figure 4:**
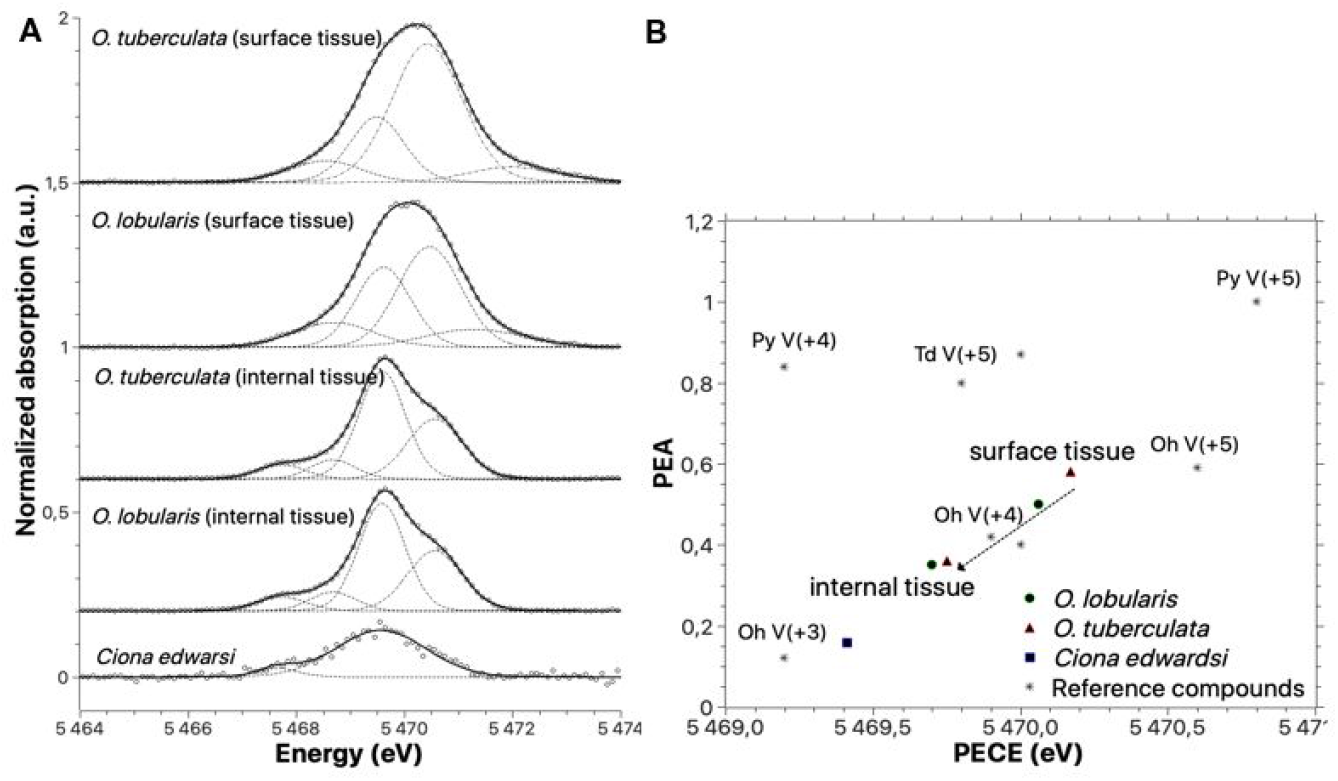
(A) Corrected pre-edge from HERFD-XANES spectra of Oscarellidae species (surface and inner tissues) and ascidian (Ciona edwardsi) collected at Maïre site (open circles). Solid lines show the fitting of the pre-edge with a series of mixed Gaussian-Lorentzian curves (sum of the 4 components in dashed lines). (B) Pre-edge features (PECE and PEA) extracted from (A). The pre-edge features are compared with those obtained for the reference compounds in which V occurs in the three oxidations states (+3, +4 or +5) in coordination geometry from octahedral (Oh) to a five-coordinate (Py) or a tetrahedral geometry (Td).

Although dominated by the vanadyl speciation, differences were observed between surface and internal tissues. HERFD-XANES pre-edge of surface tissues (*O. lobularis and O. tuberculata*) exhibits a slight shift from V (+4) features toward higher PECE and larger PEA, characteristic of V (+5) (Fig. 4.B). This suggests that a minor fraction of V (+5) can coexist with V (+4) within the sponge surface tissue. In seawater, V is mainly present as the tetrahedral vanadate ion H_n_V(+5)O_4_^(3-n)-^ (Abbasse et al., 2002; Gustafsson, 2019). The possible presence of small amounts of V (+5) at the sponge surface is probably due to the reduction of vanadate to vanadyl in the surface tissues.

On the other hand, the pre-edge features of inner tissues shift slightly toward those of V (+3) in octahedral geometry. This indicates that the average oxidation state of V within sponges changes as we go from the surface tissue (100 µm thick) to the internal tissue: Predominantly V (+4) at the surface being partly reduced to V (+3) in the inner tissue (Fig. 4.B). The case of Oscarellidae species differs from that of ascidians: In Oscarellidae, reduction to V (+3) in the internal tissues is apparently a minor process. Ascidians, assisted by Na+-dependent phosphate transporters, take up vanadate from seawater, which is then transferred to the cytoplasm and reduced to vanadyl. This reduction is done by vanabin, a vanadium-binding protein expressed in blood plasma in the presence of NADPH, glutathione, and glutathione reductase. After the reduction to vanadyl, V is further reduced to V (+3) by various, as yet unknown, mechanisms.

Finally, the HERFD-XANES spectrum of ascidian *Ciona edwardsi* exhibits pre-edge features similar to those of the reference compound, incorporating V (+3) in octahedral geometry (Fig.4). This result is in accordance with the literature. V was shown to be accumulated in ascidians mainly as a V (+3)-SO_4_ complex in a number of pioneer papers (Frank et al., 1998, 2003, 2008, 2020; Michibata, 1996; Ueki & Michibata, 2011). In these papers, V (+3) was identified as the dominating oxidation state in the blood cells, but vanadyl (+4) also contributes.

### Gene surveys evidence a convergent emergence of vanadium hyperaccumulation in Metazoa

A high-quality transcriptome and a draft genome are available for the species *O. lobularis* (Belahbib et al., 2018; Vernale et al., 2021), enabling us to search (Blast search) for genes previously reported in the literature to play a role in V transport, storage, or metabolism (in particular the genes of urochordates that were characterized in depth). None of the searched for metazoan genes that are specific to V metabolism (e.g., V-binding proteins from urochordates or V-dependant peroxidases from different protostomes, see Fig.5 and Supporting Information 5) were found in *O. lobularis*. On the other hand, gene encoding for transketolase, glutathione S-transferase and heavy metal transporting ATPase are found (reported to be involved in the V metabolic pathway of urochordates). This result is not surprising, because these genes are highly conserved across Eukaryotes. Indeed, transketolase enzymes are involved in very important metabolic pathways such as the Calvin cycle, the pentose phosphate pathway, and the oxidative stress response; glutathione S-transferase (GST) and heavy metal transporting ATPase (HMTA) are usually active in the detoxification in many species (Lee, 2008; Takahashi et al., 2012; Ueki & Michibata, 2011; Xu et al., 2016).

**Figure 5:**
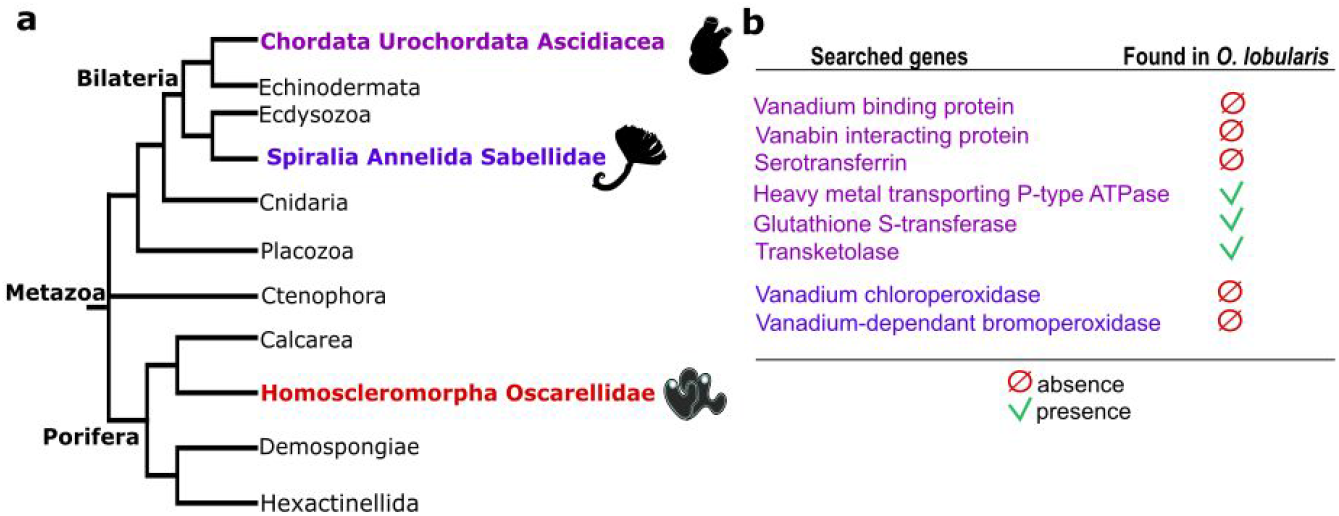
Hyperaccumulation of V emerged at least 3 times during animal evolution. (A) Phylogenetic position of the three animal lineages to date reported to perform V hyperaccumulation in the metazoan tree: Urochordata Ascidiacea, Annelida Sabellidae and Porifera Oscarellidae are very distant lineages. (B) Summary of the results of gene surveys in the genome draft of O. lobularis. Genes encoding enzymes specific to V metabolism found in urochordates (purple) and spiralians (blue) are not found in O. lobularis.

The absence of homologous vanadium specific genes argues for the independent emergence of V metabolism in the three distant metazoan lineages (Porifera Oscarellidae, Annelida Sabellidae and Urochordata Ascidiacea, Fig.5). Altogether, with the different oxidation states in which V is stored, it means that the hyperaccumulation of V in Oscarellidae relies on different, yet to be discovered, metabolic pathways from the one involved in urochordates.

### Possible involvement of Oscarella’s bacterial symbiont and functional significance?

In addition, we searched for bacterial V-dependent enzymes in the predicted proteome of *Candidatus Rhodobacter lobularis*, the most abundant bacteria associated with *O. lobularis* tissues (Jourda et al., 2015). According to the Blast search (Supporting Information 6), three sequences seem to be related to the acid phosphatase/V-dependent haloperoxidase superfamily. While two of these candidates were clearly assigned to phosphatase and N-Acetylmuramic Acid alpha-1-Phosphate (MurNAc-alpha1-P) Uridylyltransferase (data not shown), the third sequence shares a best aminoacid conservation with the active site domain of the VHPOs C-terminal region compared to acid phosphatases. Phylogenetic analyses suggest that this sequence pertains to an orthology group of unknown function, distinct from both phosphatases and vanadium haloperoxidases (VHPOs) in the PAP2 superfamily (Hemrika et al., 1997) (Supporting Information 11). Interestingly, molecular analyses have shown a conserved pattern of amino acid residues for the interaction of vanadate or phosphate in the active site of these two enzyme classes (Littlechild et al., 2002), and biochemical experiments have demonstrated that two bacterial acid phosphatases are able to bind vanadate and to exhibit a reduced halide oxidation activity (Tanaka et al., 2002).

Further sequence and biochemical analyses would be required to evaluate the functions of this protein and its possible involvement in vanadium metabolism. Even though this finding may suggest that the V metabolic pathway present in *O. lobularis* may involve at some point its symbiotic *Rhodobacter* species, there is so far no evidence that the same or a closely related bacterial species is present in *O. tuberculata* (Gloeckner et al., 2013).

Whatever the molecular mechanisms involved are, it remains to decipher what the roles are: of this V hyperaccumulation and of a V-dependent metabolism in *Oscarella* species and their associated holobiont. Several hypotheses can be proposed in the light of roles described in other species:

i. V can be used as final electron receptor for respiration by some bacteria (Antipov et al., 2000; Carpentier et al., 2005) in anaerobic conditions. This capacity may be useful if tissue oxygenation is heterogeneous in Oscarellidae, as it was shown in other sponges (Schläppy et al., 2010);
ii. In vertebrates, V was shown to promote the antioxidative response by enhancing the expression of antioxidant compounds such as glutathione and GPx (Tripathi et al., 2018). Such implication in the defense against oxidative stress might partly explain such V burden in *Oscarella* tissues. However, V is also described as a pro-oxidant and the propensity of V to act as an antioxidant or pro-oxidant depends on numerous factors (for example, concentration and oxidation state);
iii. Microbial V-dependent haloperoxidases (VHP0s) were shown to play a major role in the synthesis and diversification of secondary metabolites (Baumgartner & McKinnie, 2021; Carter-Franklin & Butler, 2004; Ishikawa et al., 2022) that are known to be numerous and powerful in sponges (Mehbub et al., 2014);
iv. In brown seaweeds, VHPOs were also shown to result in the production of volatile halocarbons (Fournier et al., 2014; Wever & Horst, 2013); but to date the profiling of odoriferous molecules secreted by sponges is only in its infancy, even though most sponge species (including the two considered species) emit a clearly perceptible unpleasant odour, which is even more intense when they are cut (Aresta et al., 2021; Christophersen et al., 1989; De Rosa et al., 2008);
v. Antifouling, anti-predation or antimicrobial properties may also come from the toxicity of the V itself (Fattorini et al., 2010; Giangrande et al., 2017; E. D. Thompson et al., 2018; Ueki & Michibata, 2011);
vi. Finally, the role of V to produce pigments, as mentioned in urochordates (Bandaranayake, 2006), may be of interest in the case of these colourful sponges (Renard et al., 2021).

### Environmental perspectives of our finding

As well as a better understanding of the bio-geochemical cycle of V, our findings open up new perspectives:

- Firstly, in view of the toxicity of V for most organisms, the *Oscarella* species may provide new clues for developing bioremediation strategies for Mediterranean marine ecosystems contaminated by V, as was already proposed for plant-based remediation strategies for continental ecosystems (Chen et al., 2021).
- Secondly, the hyperaccumulation of V by *Oscarella* species implies very specific entry and storage routes that are of particular interest for developing selective processes for the extraction of V. Indeed, in the context of climate change, there is a need to shift from traditional pyro- and hydrometallurgy processes to more environmentally friendly extraction methods. In this regard, extraction processes using bioinspired or biosourced chelators for selective recovery of the targeted metal, may offer an interesting alternative for sustainable mining (extraction in water at ambient temperature and pressure, reduced number of steps and less waste products). Although not yet developed for V, as far as we know, such strategies are currently being studied for the selective recovery of other critical metals, such as Rare Earth Elements (REEs) (Deblonde et al., 2020).

The development of bioinspired protocols requires a good knowledge of metal transfer routes. This knowledge can be obtained using omic approaches to determine molecular processes, including the nature of metabolites and proteins involved in the selective internalization of metals in living organisms. Because of the large number of studies and other resources available for *Oscarella tuberculata* and *O. lobularis* species, they would seem to be promising inspirational models.

## Acknowledgment

The authors would like to thank the following divers who assisted in collecting sponge and sediment samples: Laurent Vanbostal and Dorian Guillemain of the OSU Pythéas ; Sandrine Chenesseau and Christian Marschal of the IMBE lab ; Sandrine Ruitton of the Institut Méditerranéen d’Océanologie (MIO), and Stephane Sartoretto of the Institut Français de Recherche pour l’Exploitation de la Mer (IFREMER).

The authors would also like to thank the European Synchrotron Radiation Facility (ESRF) for providing synchrotron radiation facilities and the LA-ICP-MS platform elemental chemistry at Aix-Marseille.

For their work preparing samples and providing efficient lab facilities, the authors want to thank Geneviève Dur (CEREGE), Sandrine Chenesseau (IMBE), Alexander Ereskovsky (IMBE) and Caroline Rocher (IMBE).

Finally, thanks to Thomas Smith, native English speaker, for proofreading the manuscript.

