## Supplemental data for "Hyperaccumulation of Vanadium in animals: two sponges compete with urochordates"

**Supporting information**

Supporting Information 1: Concentrations of vanadium (mean ± standard deviation) (µg/g dw) obtained from sponge tissues and sediments collected at the different sampling sites.

| **Samples** | **Sites** | **V concentrations (µg/g dw)** |
| --- | --- | --- |
| *Oscarella lobularis* | Mejean | 18,598 ± 1,118 |
|  | Frioul | 19,431 ± 3,428 |
|  | Prado reefs | 18,306 ± 2,567 |
|  | Maïre | 15,204 ± 2,011 |
|  | Perez cave | 17,473 ± 5,392 |
|  | Riou | 19,257 ± 9,508 |
|  | Caramassaigne | 16,957 ± 1,347 |
|  | Fourmigues | 28,212 ± 6,457 |
| *Oscarella tuberculata* | Maïre | 27,009 ± 2,833 |
|  | Perez cave | 33,417 ± 88 |
|  | Caramassaigne | 31,704 ± 5,006 |
| *Aplysina cavernicola* | Maïre | 3.06 ± 1.21 |
| *Agelas oroides* | Maïre | 5.24 ± 2.49 |
| *Axinella damicornis* | Maïre | 5.08 ± 0.41 |
| *Clathrina clathrus* | Maïre | 7.21 ± 3.65 |
| *Corticium candelabrum* | Maïre | 2.3 ± 0.43 |
| *Ciona edwardsi* | Maïre | 235 ± 2.7% |
| Sediment | Maïre | 28.4 ± 5.8% |
|  | Frioul | 3.82 ± 18% |
|  | Prado reefs | 30.1 ± 3,8% |
|  | Perez cave | 5.89 ± 12% |
|  | Caramassaigne | 4.76 ± 25.5% |
|  | Riou | 2.23 ± 23% |
|  | Mejean | 23.4 ± 5.9% |
|  | Fourmigues | 16.0 |

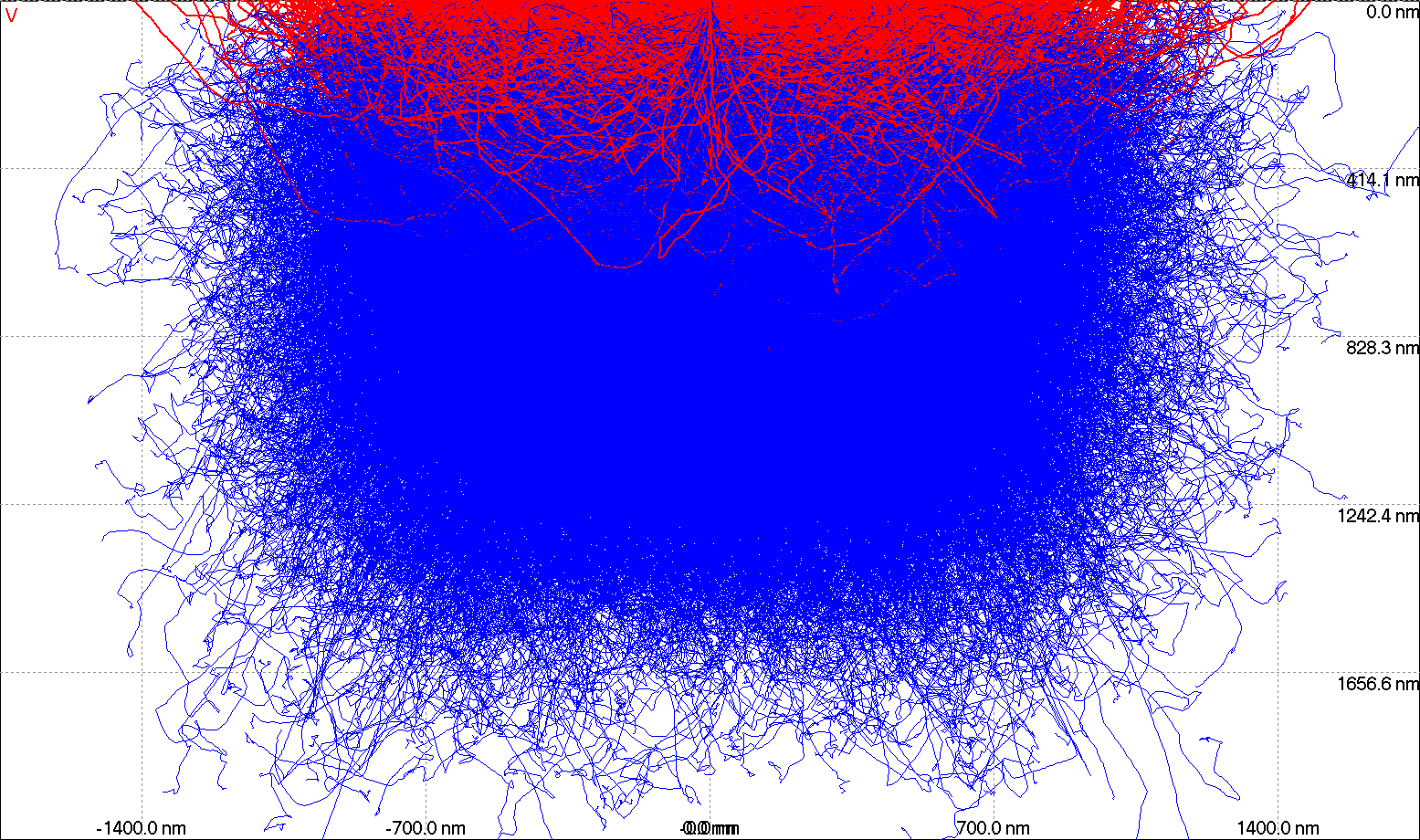

Supporting Information 2: Monte Carlo simulation of electrons trajectories in the pieces of marine sponges containing vanadium. The blue trajectories represent the incident electrons and the red trajectories line represent the back-scattered electrons.

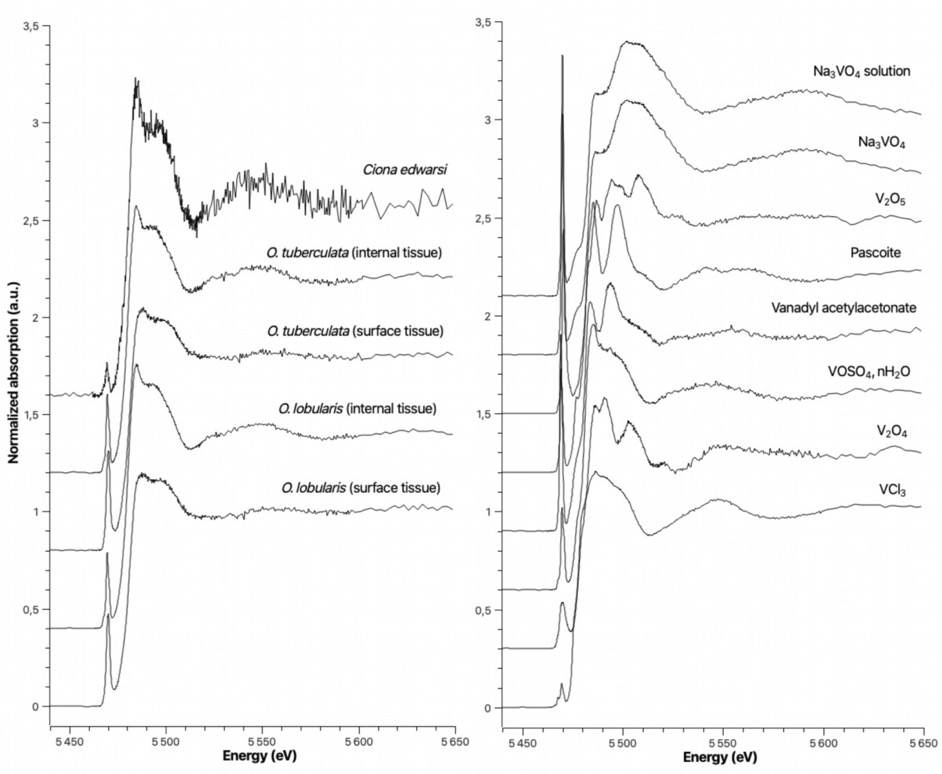

**A**

**B**

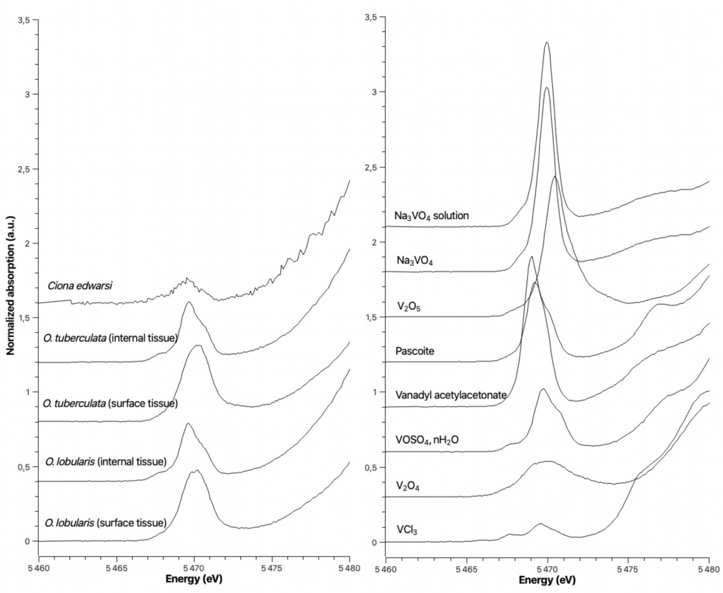

Supporting Information 3. Experimental V K-edge HERFD-XANES spectra of (A) of Oscarellidae species (surface and internal tissues) and ascidian (Ciona edwardsi) collected at Maïre site and of (B) reference compounds. A zoom on the pre-edge region is also provided.  Dataset of reference compounds is available in the SSHADE database (Chaurand, Perrine; Couturier, Julien; Levard, Clément; De Pao Mendonca, Kassandra (2021): V K edge XAS HERFD of V reference compounds 10K. SSHADE/FAME (OSUG Data Center). Dataset/Spectral Data. doi:10.26302/SSHADE/EXPERIMENT_PC_20220301_001)

*Supporting Information 4.* *Pre-edge features (PECE, PEA) obtained by fitting a baseline-corrected pre-edge peak with a series of mixed Gaussian-Lorentzian curves (pseudo-Voigt functions).*

|  | PECE* (eV) | PEA** | Component 1 | | Component 2 | | Component 3 | | Component 4 | |
| --- | --- | --- | --- | --- | --- | --- | --- | --- | --- | --- |
| **Sample (Maïre site)** |  |  | Center (eV) | Area | Center (eV) | Area | Center (eV) | Area | Center (eV) | Area |
| *Oscarella lobularis*  Surface tissue | 5,470.0 | 0.50 | 5,468.7 | 0.14 | 5,469.6 | 0.31 | 5,470.4 | 0.45 | 5,471.2 | 0.12 |
| *Oscarella lobularis*  Inner tissue | 5,469.7 | 0.35 | 5,467.7 | 0.05 | 5,468.7 | 0.06 | 5,469.6 | 0.36 | 5,470.6 | 0.24 |
| *Oscarella tuberculata*  surface tissue | 5,470.1 | 0.58 | 5,468.6 | 0.11 | 5,469.5 | 0.26 | 5,470.4 | 0.71 | 5,472.0 | 0.1 |
| *Oscarella tuberculata*  Inner tissue | 5,469.7 | 0.36 | 5,467.8 | 0.05 | 5,468.6 | 0.02 | 5,469.6 | 0.46 | 5,470.7 | 0.21 |
| *Ciona edwardsi* | 5,469.4 | 0.16 | 5,467.7 | 0.03 | 5,469.6 | 0.30 |  |  |  |  |
| **Reference compounds** |  |  |  |  |  |  |  |  |  |  |
| VCl_3_  Oh V (+3) | 5,469.2 | 0.12 | 5,465.9 | 0.01 | 5,467.8 | 0.06 | 5,469.6 | 0.15 | 5,470.6 | 0.03 |
| V_2_O_4_  Oh V (+4) | 5,470.0 | 0.40 | 5,470.0 | 0.81 |  |  |  |  |  |  |
| VOSO_4_, nH_2_O  Oh V (+4) | 5,469.9 | 0.42 | 5,467.8 | 0.05 | 5,468.7 | 0.04 | 5,469.7 | 0.45 | 5,470.7 | 0.33 |
| Vanadyl acetylac.  Py V (+4) | 5,469.2 | 0.84 | 5,468.5 | 0.29 | 5,468.9 | 0.57 | 5,469.7 | 0.85 |  |  |
| Pascoit***  Oh V (+5) | 5,470.6 | 0.59 | 5,468.0 | 0.06 | 5,469.2 | 0.52 | 5,470.1 | 0.41 |  |  |
| V_2_O_5_  Py V (+5) | 5,470.8 | 1.00 | 5,470.4 | 0.51 | 5,470.9 | 1.44 | 5,473.7 | 0.08 | 5,467.9 | 0.02 |
| Na_3_VO_4_  Td V (+5) | 5,470.0 | 0.87 | 5,469.0 | 0.25 | 5,470.0 | 1.14 | 5,470.7 | 0.39 |  |  |
| Na_3_VO_4_ solution  Td V (+5) | 5,469.8 | 0.80 | 5,469.1 | 0.28 | 5,470.0 | 1.32 | 5,471.2 | 0.04 |  |  |

** PECE: pre-edge peak centroid energy, area-weighted average of the position in energy of each component, **PEA: pre-edge peak area, sum of the integrated area of each component normalized to V_2_O_5_ pre-edge area.*

*Geometry: Oh, octahedral (6-coordinate); Td, tetrahedral (4-coordinate); Py, square pyramidal (5-coordinate).*

**** from (Chaurand et al., 2007).*

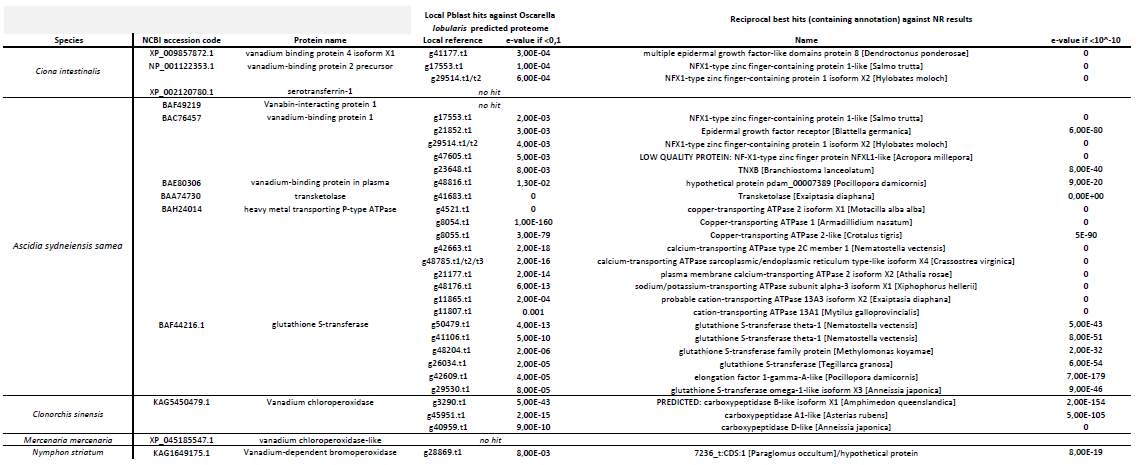

Supporting Information 5: List of query sequences used for Blast P against the predicted proteome of Oscarella lobularis and of reciprocal best hits against NR database.

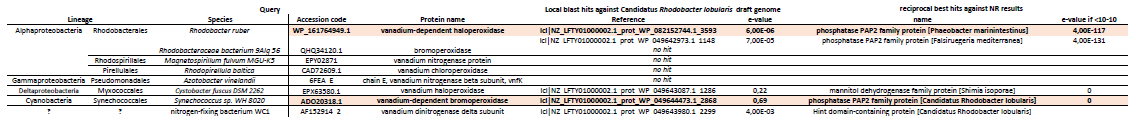

Supporting Information 6: List of query sequences used for Blast search against the genome of Candidatus Rhodobacter lobularis and of the reciprocal best hits obtained against the NR database.

Supporting Information 7: Bioconcentration factors (BCF) calculated for each species at Maïre site.

| **Species** | **BCFs (µg/g)** |
| --- | --- |
| ***Oscarella lobularis*** | **535** |
| ***Oscarella tuberculata*** | **951** |
| *Aplysina cavernicola* | 0.11 |
| *Agelas oroides* | 0.18 |
| *Axinella damicornis* | 0.18 |
| *Clathrina clathrus* | 0.25 |
| *Corticium candelabrum* | 0.08 |
| *Ciona edwardsy* | 8.27 |

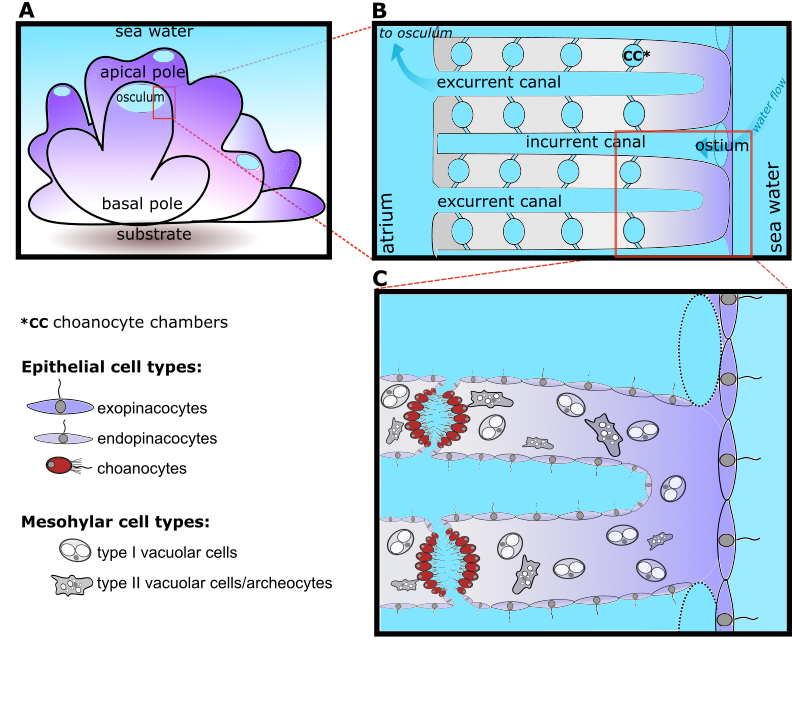

Supporting Information 8: General organisation of Oscarella lobularis and position of cell types.

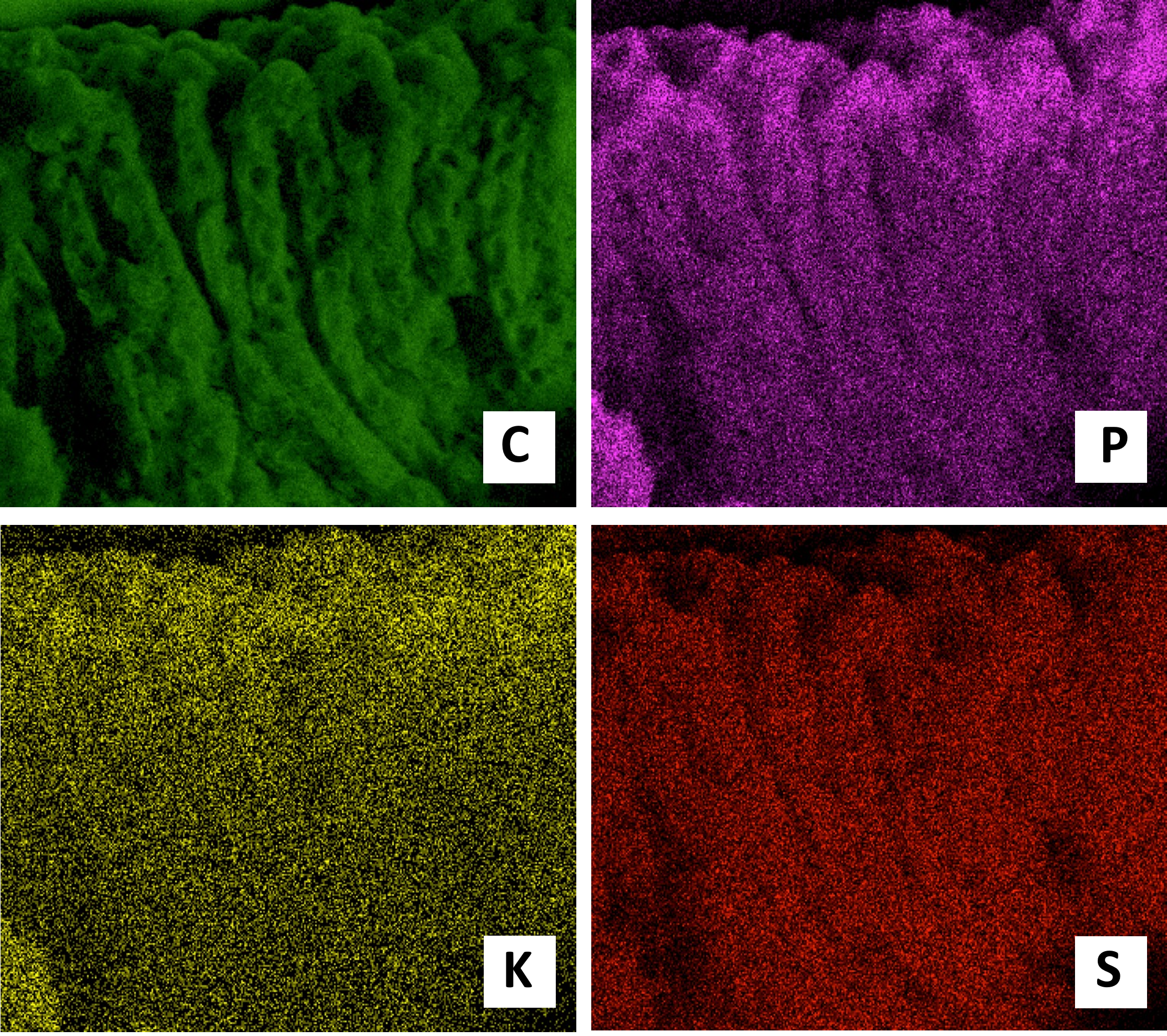

Supporting Information 9: EDX map of essential elements like carbon (C), phosphorus (P), potassium (K), and sulphur (S) in Oscarella lobularis tissues. Carbon, sulfur and potassium have a homogeneous distribution within the tissues. The difference in signal intensity between the surface and the interior of the body indicates a higher concentration of phosphorus on the surface.

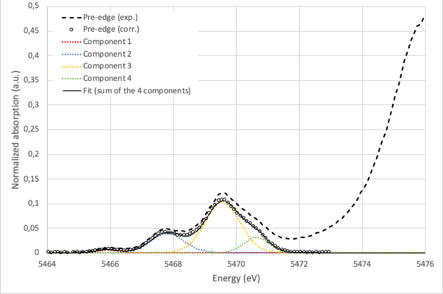

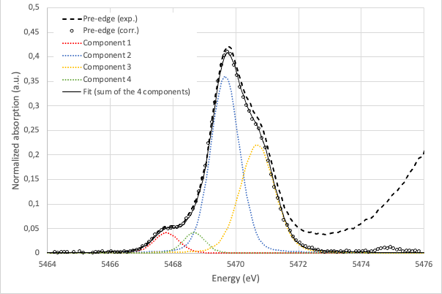

**B**

**A**

*Supporting Information 10. Examples of fitting of the corrected pre-edge peak obtained for (A) VCl_3_ (solid), and (B) VOSO_4_, nH_2_O with a series of mixed Gaussian-Lorentzian curves. The sum of the integrated area of the 4 components gives the area of the pre-edge peak (PEA in Table Supporting Information 4) and the area-weighted average of the positions of the 4 components gives the centroid position of the pre-edge peak (PECE in Table Supporting Information 4).*

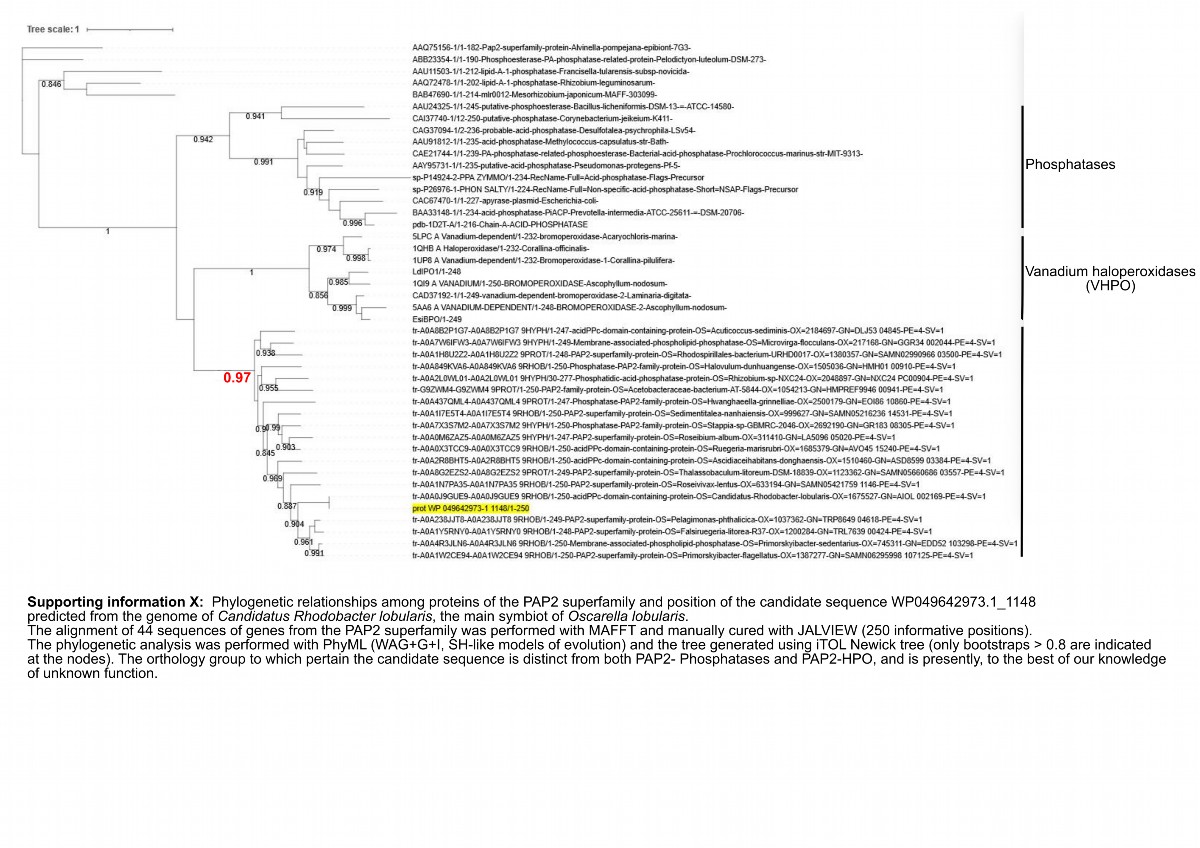

Supporting information 11: Phylogenetic relationships among proteins of the PAP2 superfamily and position of the candidate sequence WP049642973.1_1148 predicted from the genome of Candidatus Rhodobacter lobularis, the main symbiot of Oscarella lobularis. The alignment of 44 sequences of genes from the PAP2 superfamily was performed with MAFFT and manually cured with JALVIEW (250 informative positions). The phylogenetic analysis was performed with PhyML (WAG+G+I, SH-like models of evolution) and the tree generated using iTOL Newick tree (only bootstraps > 0.8 are indicated at the nodes). The orthology group to which pertain the candidate sequence is distinct from both PAP2- Phosphatases and PAP2-HPO, and is presently, to the best of our knowledge of unknown function.
